# Deep learning-based segmentation and quantification of podocyte foot process morphology

**DOI:** 10.1101/2021.06.14.448284

**Authors:** Linus Butt, David Unnersjö-Jess, Martin Höhne, German Sergei, Anna Witasp, Annika Wernerson, Jaakko Patrakka, Peter F. Hoyer, Hans Blom, Bernhard Schermer, Katarzyna Bozek, Thomas Benzing

## Abstract

The kidneys constantly filter enormous amounts of fluid, with almost complete retention of albumin and other macromolecules in the plasma. Diseases of podocytes at the kidney filtration barrier reduce the intrinsic permeability of the capillary wall resulting in albuminuria. However, direct quantitative assessment of the underlying morphological changes has previously not been possible. Here we developed a deep learning-based approach for segmentation of foot processes in images acquired with optical microscopy. Our method – Automatic Morphological Analysis of Podocytes (AMAP) – accurately segments foot processes and robustly quantifies their morphology. It also robustly determined morphometric parameters, at a Pearson correlation of r > 0.71 with a previously published semi-automated approach, across a large set of mouse tissue samples. The artificial intelligence algorithm wasWe applied the analysis to a set of human kidney disease conditions allowing comprehensive quantification of various underlying morphometric parameters. These results confirmed that when podocytes are injured, they take on a more simplified architecture and the slit diaphragm length is significantly shortened, resulting in a reduction in the filtration slit area and a loss of the buttress force of podocytes which increases the permeability of the glomerular basement membrane to albumin.

## INTRODUCTION

The capacity of the mammalian kidney to filter vast amounts of fluid (180 L/day in humans) and to almost completely restrict the passage of macromolecules (e.g. albumin) relies on the intricate glomerular filtration barrier. This barrier consists of a fenestrated endothelium, the glomerular basement membrane (GBM) and specialized post-mitotic epithelial cells, called podocytes^1^. Damage to any of the three layers results in an increased filter permeability, with albuminuria being the most prominent clinical symptom^1^. Due to the nanoscale dimensions of the filter, electron microscopy is widely used in clinical pathology and research to assess morphological alterations associated with glomerular injury. Pathological alterations of podocytes, particularly affecting their so-called foot processes (FPs) and the slit diaphragm (SD) as their only cell junction, are seen in most types of glomerular diseases. These alterations, referred to as *foot process effacement*, include widening and ultimately loss of FPs together with a progressive shortening of total SD length^2^. Super-resolution light microscopy techniques allow for visualizing and quantifying morphological alterations upon FP effacement^3–6^. Recently, our group has used this approach in a mouse model of hereditary focal and segmental glomerulosclerosis (FSGS) to provide a model of kidney ultrafiltration in which size selectivity of the filtration barrier is dependent on GBM compression^7^. In that study, FP morphology correlated robustly with levels of albuminuria in mutant mice. Integrating these data in a mathematical model for ultrafiltration, it was proposed that injured podocytes fail to adequately counteract filtration pressure, leading to a relaxed matrix within the GBM resulting in increased permeability to albumin. Until now, the aforementioned morphological analyses had to be carried out manually or semi-automatically, which is not only time consuming but also investigator-dependent, currently impeding their broad use in research and diagnostics.

Here, we combined previously established morphological analyses with a machine learning algorithm to enable fully automated segmentation and quantification of podocyte ultrastructure. New machine learning methods automate bioimage analysis at a human-level accuracy with cancer histopathology as one of its most prominent applications^8,9^. In nephrology, deep learning (DL) methods have been proposed for the segmentation of entire renal structures or entire podocytes^10–12^. However, none of the existing segmentation methods allows recognition of podocyte substructures.

Here we propose Automatic Morphological Analysis of Podocytes (AMAP) – a fully automated method for segmentation of FPs and the SD from optical microscopy images of podocytes. Our method is based on DL instance segmentation and is trained and tested on a broad range of disease and imaging conditions, correctly detecting FPs across our datasets and reproducing morphometric parameters originally extracted using an ImageJ macro – referred to as macro throughout the manuscript. It successfully generalizes to human samples and to images acquired using confocal microscopy, a system commonly available in clinical pathology labs. Our approach opens up possibilities for systematic studies of nanoscale pathologies occurring in kidney disease.

## RESULTS AND DISCUSSION

Using a previously published STED imaging protocol^3^, we generated 209 images representing healthy and diseased mouse tissue at various ages and degrees of podocyte effacement. The images represent FP shapes by staining the SD protein nephrin (see Fig. 1a for an overview). Annotation of these images, including both SD and FP areas (Supplementary Fig. 1), was generated with a network pre-trained on a small set of macro-labeled images. These annotations were further manually corrected to obtain a possibly complete annotation of each image. For improved accuracy each file was annotated by at least two individuals.

**Figure 1.**
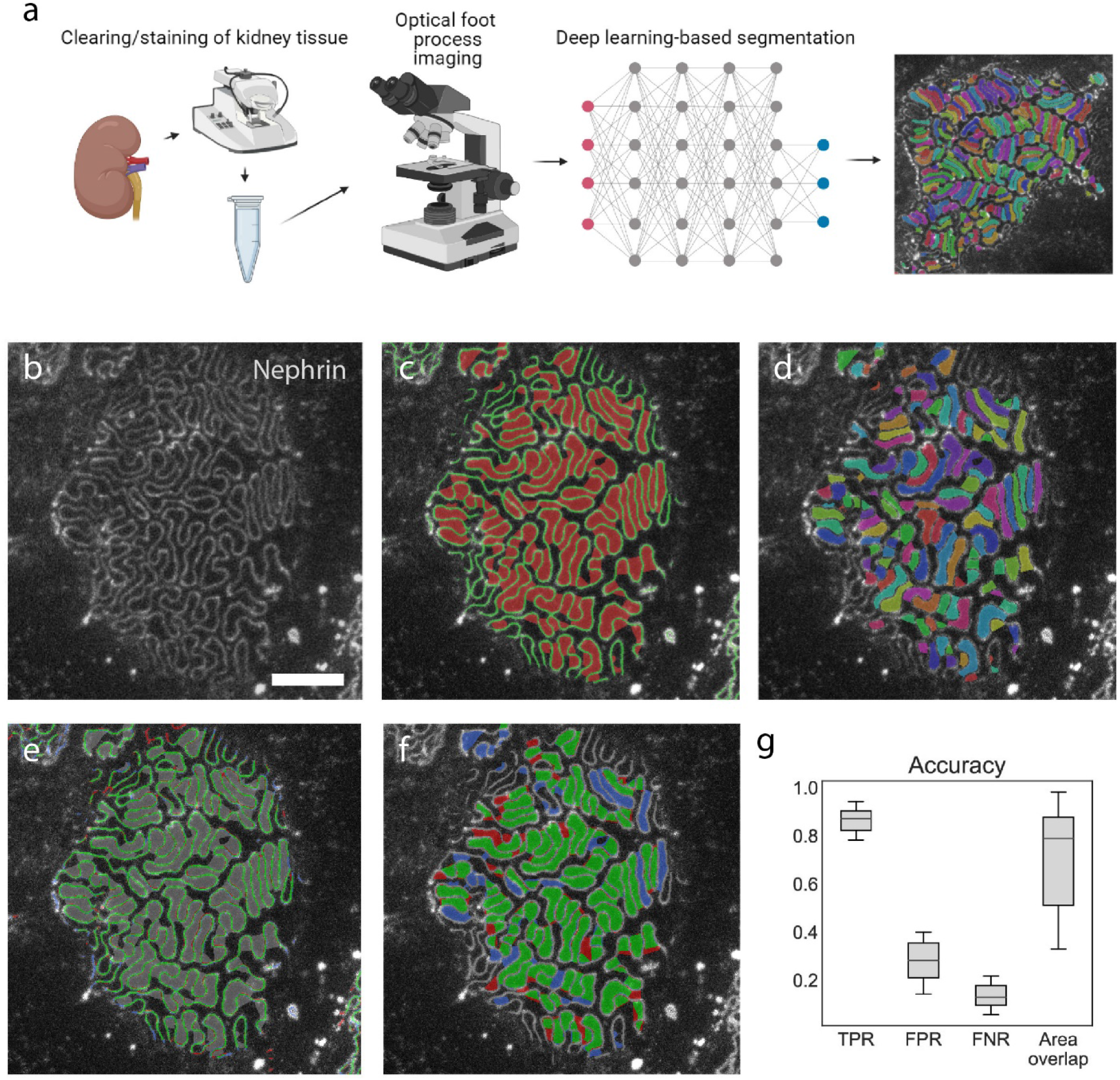
Overview of AMAP. (a) Schematic overview. Previously published sample preparation and imaging protocols are used for visualization of FPs. Convolutional neural network is applied to extract FP and SD regions allowing for fully automatized segmentation of FPs in kidney samples. (b) Nephrin-stained STED image from the test set. Scale bar 2 µm. (c) Outcome of semantic segmentation. FP pixels are marked in red, SD pixels in green. (d) Outcome of instance segmentation. Separate FP instances are marked with different colors. (e) Accuracy of SD semantic segmentation. Pixels correctly predicted as SD are marked in green, those labeled but not predicted as SD (false negatives) are marked in red, those predicted but not labeled as SD (false positives) are marked in blue. (f) Same as (e) for FP pixels. (g) FP instance prediction accuracy in the entire test set of 209 pictures. True positive rate (TPR), false-positive rate (FPR), false-negative rate (FNR) are quantified relative to the number of manually determined FPs in a given image. Area overlap is quantified as intersection over union (IoU). FPR includes a large proportion of not labeled FPs and does not accurately reflect the actual errors in prediction.

We next trained a segmentation convolutional neural network (CNN) on the training set, incorporating both semantic and instance segmentation (Fig. 1b-d)^13^. Semantic segmentation classifies pixels into three classes: background, FP, and SD (Fig. 1c). Instance segmentation additionally assigns FP pixels into separate FP instances (Fig. 1d, Supplementary Fig. 2).

We trained the network and inspected its segmentation performance on the test set. On average, AMAP assigned correct classes to 73% and 76% FP and SD pixels, respectively. Relative to the number of pixels in the FP and SD class, 23-27% of pixels were misclassified, i.e., incorrectly assigned either to background, FP, or SD. Based on visual inspection, we noted that these errors were sometimes due to incomplete annotation and uncertainty in the exact FP and SD boundaries (Fig. 1e-f).

The FP pixels were additionally split into separate instances, and we matched predicted instances with labels. Predicted instances overlapping with a labeled FP were counted as true positives and area of overlap with the matched label was quantified. In test images, we detected 87% of all labeled FPs (Fig. 1g). Matched FP instances overlapped on average at 0.74 quantified as Intersection over Union (IoU) of labeled and predicted instances. Relative to the number of labeled FPs, 28% more FPs were predicted with AMAP, while 13% were not detected (Fig. 1g). Notably, some potential false positive detections were due to missing labels and should not be considered errors.

As further validation, we asked if AMAP allows to accurately reproduce morphometric parameters extracted using a previously published macro-based analysis of the podocin^R231Q/A286V^ FSGS mouse model and their control littermates^7^. Podocin^R231Q/A286V^ mice showed quantifiable morphological alterations of individual FPs as well as the SD pattern that correlated with albuminuria. We ran AMAP on 174 nephrin-stained images used in the original publication (Fig. 2a), including control and mutant mice of different ages. Out of 19,676 labeled FPs in the published dataset, we correctly detected 95% (Fig. 2b) with an area overlap of 0.81 IoU on average. However, we detected 25,193 additional FPs not detected in the published dataset. Visual inspection of these predictions suggests that they predominantly represent FPs which were overlooked in the macro analysis. Our method failed to detect 5% of all annotated FPs.

**Figure 2.**
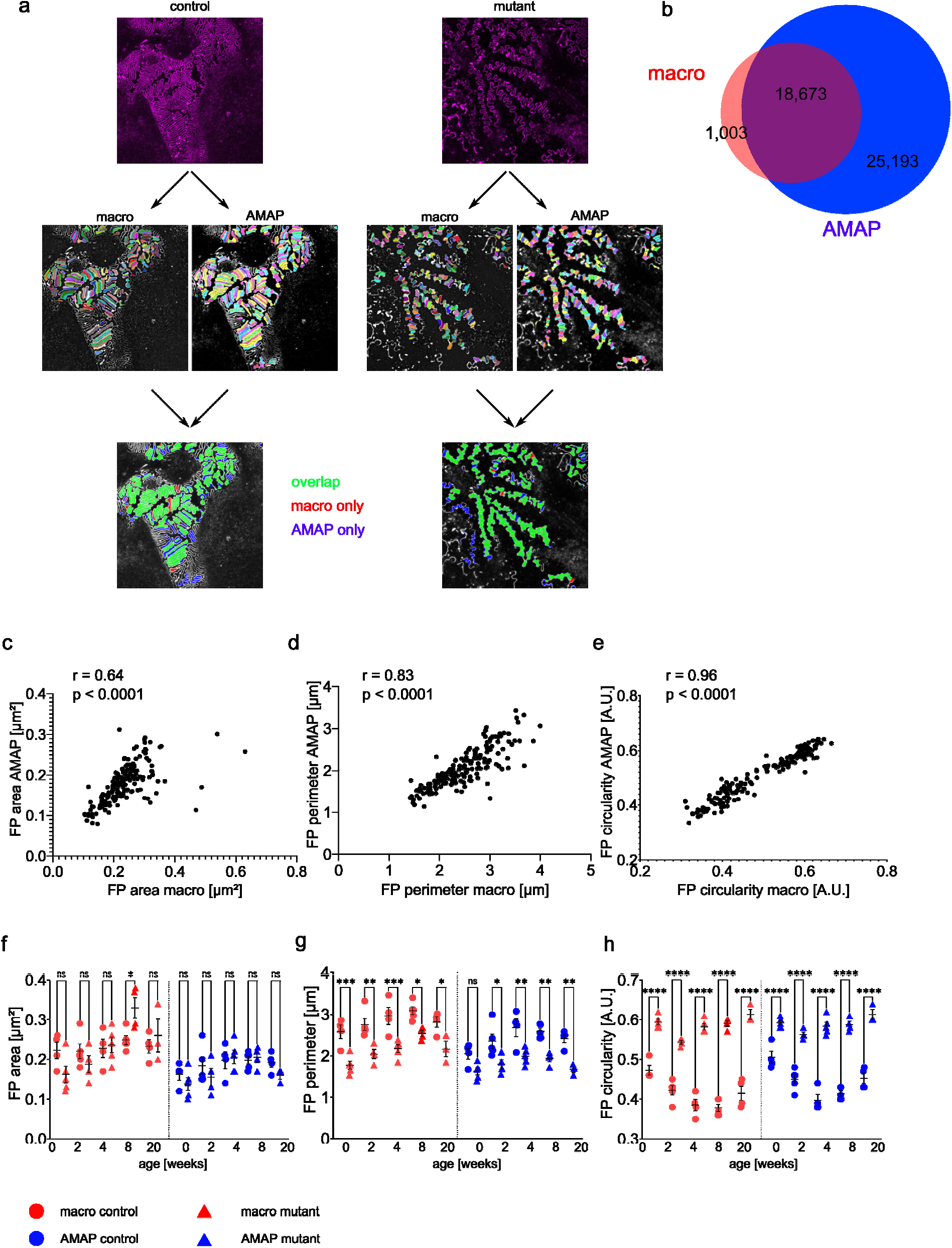
Comparison of AMAP- and macro-based FP and SD morphology in podocin^R231Q/A286V^ (“mutant”) and control mice from a published dataset. (a) Illustration of macro and AMAP FP segmentation in a control and a mutant mouse. Differences and overlap of detection are color-coded as indicated. (b) Venn diagram showing the overlap and differences in FP detection. (c)-(e) Correlation of FP area (c), FP perimeter (d) and FP circularity (e) between the macro- and AMAP-assigned FPs. Measurements of all three parameters correlate significantly between the approaches. Each dot represents the values originating from one image (n = 174 images). r = Pearson correlation coefficient, p = p-value. (f)-(h) Comparison of the values for FP area (f), FP perimeter (g) and FP circularity (h) in macro-assigned (red) or AMAP-assigned (blue) FPs in control and age-matched mutant mice. With the exception of FP area at 8 weeks and FP perimeter at 0 weeks, the detection of significant differences is comparable between the approaches. Each dot/triangle represents one mouse. Data presented as mean ± SEM. Sidak’s multiple comparison test was performed to determine statistical significance. * p < 0.05, ** p < 0.01, *** p < 0.001, **** p < 0.0001.

Instead of manually verifying the over 40,000 FPs segmented by AMAP, we inspected to what extent AMAP reproduces the morphometry of labeled FPs at each age and disease stage (See Supplementary Fig. 3 for an overview of quantified parameters). We found an overall high agreement of the morphometric parameters (Fig. 2c-e). Values for area and perimeter inferred by AMAP were lower compared to the macro-extracted ones (Fig. 2f-g). However, morphological parameters of FPs detected both by AMAP and macro show high agreement (r > 0.85, Supplementary Fig. 4), suggesting that the additional FPs detected by AMAP tend to be smaller. Consistent with earlier findings, perimeter and circularity extracted by AMAP allowed distinguishing control from mutant mice, whereas FP area alone is insufficient to make this distinction (Fig. 2f-h). These results illustrate that AMAP robustly reflects alterations to FP morphology characteristic for the progression of FSGS.

It has previously been shown that FP effacement affects not only the morphology of individual FPs but also the overall SD length^3,6,7^. Additionally, SD length has been linked to the occurrence of albuminuria^7^. Therefore, we developed a method to automatically assign regions of interest (ROIs) based on the SD staining and quantify the SD length within these ROIs (Fig. 3a, Supplementary Fig. 3). AMAP resulted in 20 % lower values for SD length per area compared to the macro (Supplementary Fig. 5a), owing to the 25 % on average larger ROIs assigned by AMAP (Fig. 3 b, Supplementary Fig. 5b), whereas the total SD length quantified by AMAP was only 5 % larger (Fig. 3c). Even though there was a difference in absolute values of ROI area and total SD length detected, the respective values correlated robustly between the macro and AMAP (Fig. 3d-e). Consequently, the previously observed decrease of SD length per area in mutant as compared to control mice was reliably detected using AMAP (Fig. 3f-g). Notably, AMAP-derived values also allow for delineating the decrease in SD abundance with aging (Supplementary Fig. 5c). Taken together, AMAP shows the potential to detect pathological alterations in individual FP morphology as well as in global SD abundance. Furthermore, plotting the mean values of AMAP-derived SD length per area against the levels of albuminuria reveals a similarly robust correlation between SD length and albuminuria as was already shown for the macro-based SD length values (Fig. 3 h).

**Figure 3.**
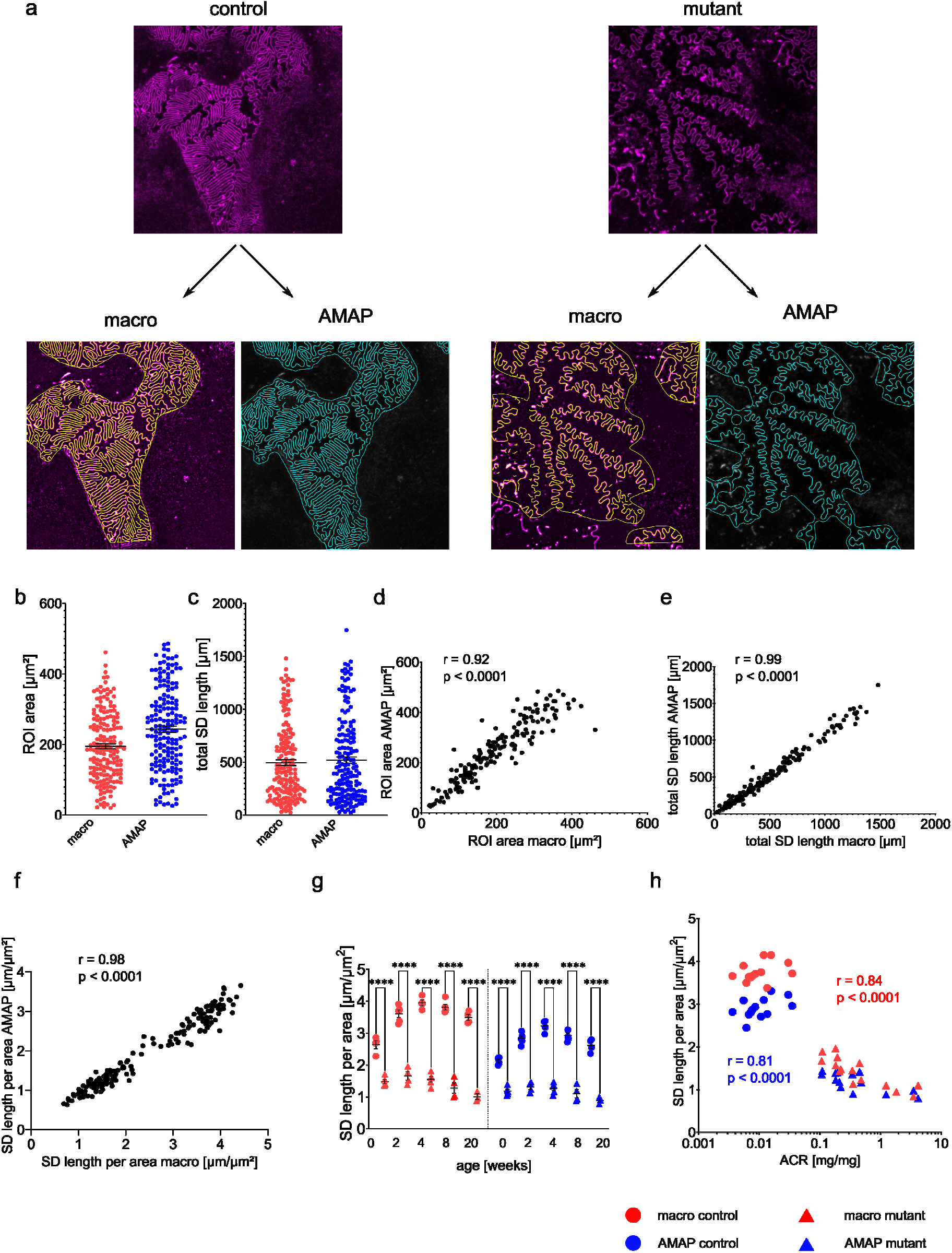
Comparison of SD length in podocin^R231Q/A286V^ (“mutant”) and control mice calculated with the macro and AMAP. (a) Segmentation of the SD length and ROIs with the two approaches in control and a mutant mouse. (b) Area of manually assigned (red) and AMAP-assigned (blue) ROIs. Each dot represents the ROI area of one image (n = 174 images). Horizontal bars represent the median. (c) Total SD length quantified with the macro (red) and AMAP (blue) ROIs. Each dot represents the ROI area of one image (n = 174 images). Horizontal bars represent the median. (d)-(f) Correlation of ROI area (d), total SD length (e) and SD length per area (f) of macro- and AMAP-assigned ROIs. Each dot represents the ROI area of one image (n = 174 images). r = Pearson correlation coefficient, p = p-value. (g) Comparison of the SD per area values in macro- (red) or AMAP-assigned (blue) ROIs in control and age-matched mutant mice. Differences between genotypes are equally detected between the two approaches. Each dot/triangle represents one mouse. Data are presented as mean ± SEM. Sidak’s multiple comparison test was performed to determine statistical significance. **** p < 0.0001. (h) Scatter plots of the mean SD length per area (macro in red, AMAP in blue) against the urinary albumin creatinine ratio (ACR). Two-tailed Spearman’s rank correlation was used to determine statistical significance. r, Spearman’s rank coefficient.

The imaging protocol we have utilized in the above analyses is lengthy (3-4 days) and requires the use of super-resolved microscopy. We have recently published a protocol for fast and straightforward confocal microscopy-based imaging of FPs (referred to as *fast protocol* throughout the manuscript), which reduces the time needed for sample preparation and imaging to only 5 h^4^. To adapt the method to the lower optical resolution of this protocol we generated an additional set of 15 and 40 annotated images of human and mouse tissue, respectively, obtained with the fast protocol. To facilitate the annotation of mouse tissue, we also generated matched images using STED microscopy. We inferred FP and SD segmentations using the network described above on STED images, corrected them manually, and used the results as labels of the confocal images (Supplemental Fig. 6a). Images of human tissue were annotated manually (Supplementary Fig. 6b). We added the 55 images to the existing training set and ran another training procedure on the network initially trained on the STED microscopy images. As evaluated by visual inspection, this retraining resulted in improved segmentation results also for images of lower quality and resolution (Supplementary Fig. 6c).

We tested the adapted segmentation and analysis workflow on a set of 43 confocal microscopy images of human tissue. Most of the images, except for the FSGS patient, are part of a previously published set of images^4^, including congenital nephrotic syndrome with mutations in the *TRPC6/NPHS2* genes, minimal change disease (MCD), FSGS, and IgA nephropathy.

Based on visual inspection, AMAP offers good accuracy in the more challenging images of human tissue imaged using the fast protocol (Fig. 4a). Moreover, morphometric parameters of the segmented FPs reflected different degrees of effacement and morphological changes in each patient in agreement with what has been previously published^4^ (Fig. 4b-c). In the FSGS mouse model (see Figures 2 and 3), a decrease in SD length is always accompanied by an increase in FP circularity. In contrast, the FSGS patient showed a relatively high SD length and a high circularity as compared to the other diagnoses, whereas the opposite was observed for the *TRPC6/NPHS2* patient. We therefore visualized multiple morphometric parameters (SD length, SD grid index, FP area, FP perimeter, FP circularity (Supplementary Fig. 3)) using umap^14^ (Fig. 4d), which illustrates morphometry-based grouping among the different diagnoses. For example, the FSGS and the MCD patients appear away from one another in the plot. Both of these patients show similar SD lengths (Fig. 4b) but a significantly different circularity (Fig. 4c). The *TRPC6/NPHS2* patient is positioned away from control samples and shows a more extensive spread of data points, reflecting the more heterogeneous effacement pattern in this patient as compared to the others. To show the throughput capacity of AMAP we imaged large field-of-views of 155*155 µm^2^ (taking around 2-6 minutes to acquire with a high-end confocal microscope). AMAP segmented 6,452 and 176 FPs from the control and *TRPC6/NPHS2* patients, respectively, in < 1 h (Fig. 4 e-f) (would require ∼ 7 h of hands-on work for a trained user). While based on a limited number of tissue samples, these results suggest that our fully automated approach for segmentation and morphometric parameter quantification opens up possibilities for a comprehensive, large-scale description of FP effacement across kidney pathologies.

**Figure 4.**
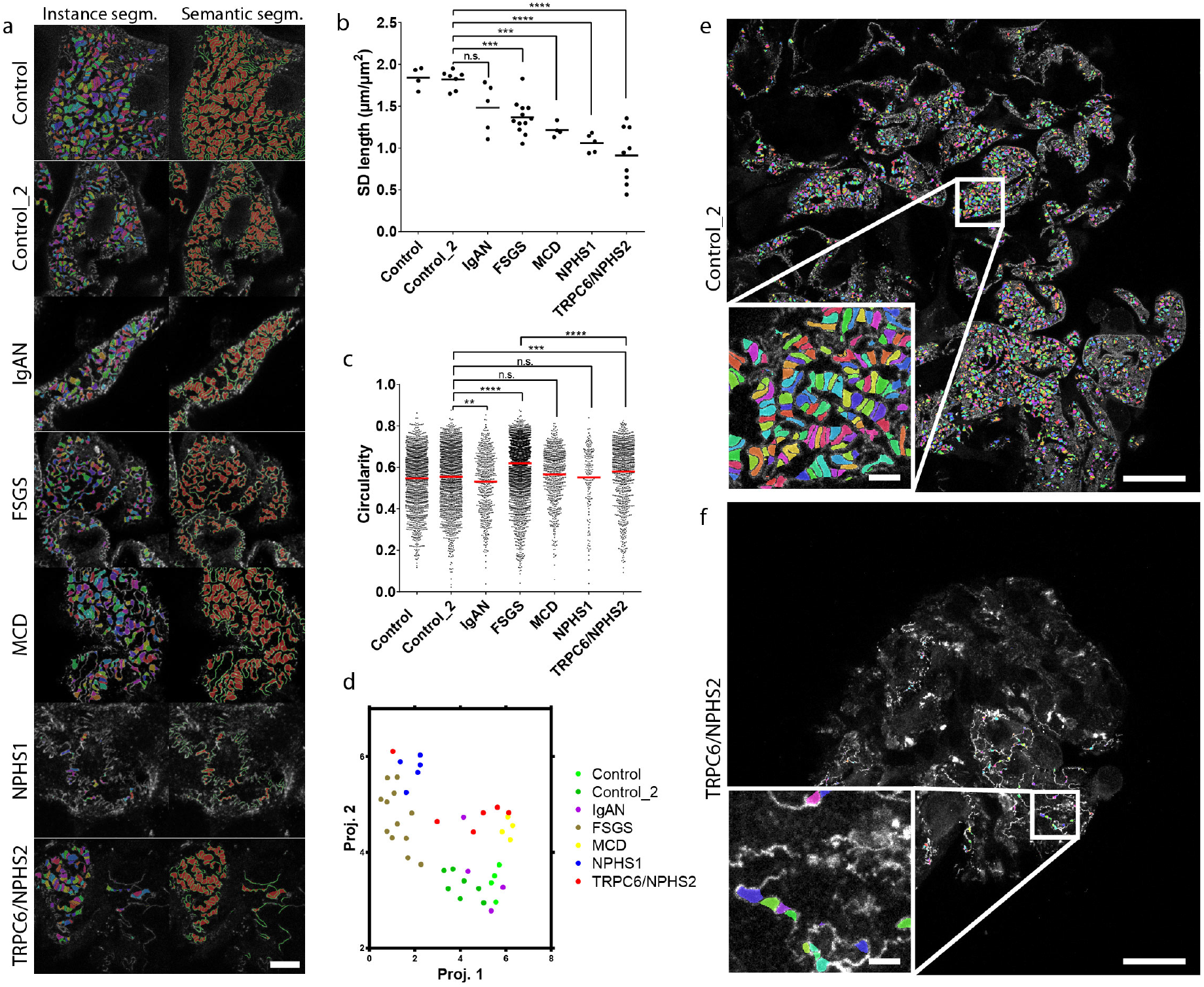
AMAP-based FP morphometry in human patients. (a) Each row represents one patient as indicated with raw images overlaid with the results of instance (left) and semantic (right) segmentation. All samples were stained for nephrin except for the NPHS1 patient which was stained for podocin. Scale bar 5 µm. (b) SD length per area for all patients shows a significant decrease for all patients except for the patient with IgAN. Each dot represents one image (one patient per diagnosis). The black line represents the mean. Tukey’s multiple comparison test was performed to determine statistical significance. *** p < 0.001, **** p < 0.0001. (c) Circularity score of FPs from all patients. Each data point represents one FP. The red line represents the mean. Dunnet’s multiple comparison test was performed to determine statistical significance. ** p < 0.01, *** p < 0.001, **** p < 0.0001. (d) Multiparameter projection (umap) of all patients. (e-f) 155 um field-of views maximum intensity projections of z-stacks of a control patient (a) and a sclerosed glomerulus of the *TRPC6/NPHS2* patient. Insets show zoomed views of the indicated areas. 6,452 FPs are segmented from the image in (e), showing the data throughput capacity of AMAP. Scale bars 20 µm and 2 µm (inset).

In recent years, light microscopy-based imaging of podocyte ultrastructure has gained an enormous momentum. Not only does this technique complement electron microscopy imaging, but also offers the possibility of efficiently imaging much larger tissue areas and enables quantitative analysis of pathological alterations in disease progression^3,6,7^. Combining quantitative analyses with biophysical modelling of ultrafiltration, our group has previously proposed a model, which mechanistically linked FP effacement with the occurrence of albuminuria^7^. Although powerful, these quantitative analyses still rely on manual assignment of ROIs and a manual step in the segmentation of individual FPs. This is not only time-consuming but also potentially prone to bias.

By utilizing DL, we present a fully automatic segmentation of FPs and the SD. We demonstrate that this approach successfully reproduces previously published results in a mouse model for FSGS^7^. Thus, AMAP can be applied to quantify alterations to FP morphology with the same accuracy and substantially higher throughput as our previous approach, while eliminating user bias. Importantly, AMAP confirmed the correlation of SD length and levels of albuminuria, thereby supporting the previously proposed mechanism of albuminuria upon podocyte injury^7^.

We further validate that the DL network can be adapted to segment the SD and FPs in human patient samples. These images were acquired using a recently published fast and simple protocol utilizing confocal microscopy^4^. Interestingly, we show that multiparameter analysis of FP morphometry in these patients shows distinct groupings for different disease types. This finding indicates that the view of FP effacement as a uniform process might have to be revised and that effacement patterns could differ depending on the underlying disease. Even though the presented dataset is limited and that more data points are needed to establish these findings in the future, we still suggest that AMAP could lead to a better understanding of the link between FP morphology and disease progression. Depending on the data size, the method can be deployed on a local machine or any more advanced computer station equipped with a graphics processing unit (GPU) which substantially speeds up the CNN processing. Apart from saving working hours, incorporating AMAP in clinical renal pathology routines could allow for more precise diagnostics due to the additional morphometric information it provides.

In summary, AMAP, the first imaging, and analysis strategy allowing for non-biased, fully automated quantification of FP morphology at the nanoscale, confirms our previous finding that the SD length correlates robustly with levels of albuminuria in a mouse model of hereditary FSGS, thereby supporting the mechanistic link between a simplified podocytes ultrastructure and the occurrence of albuminuria. We demonstrate that AMAP is readily applicable to human samples processed with our recently published fast protocol, which no longer requires sophisticated super-resolution microscopy. The combination of AMAP and the fast protocol might in the future allow for multi-scale three-dimensional and quantitative kidney diagnostics using only one sample preparation, imaging and analysis workflow^10–12^.

## MATERIALS AND METHODS

All methods can be found in the supplement.

## ACKNOWLEDGEMENTS

We thank the CECAD Imaging Facility, Cologne, Germany and the Advanced Light Microscopy (ALM) Facility, Solna, Sweden for their support in the acquisition of microscopy data. This work was supported by the Clinical Research Unit (CRU) 329 (KFO 329; A1, A6 and A7) as well as partly by FOR 2743 of the Deutsche Forschungsgemeinschaft (DFG). KB and GS were supported by the North Rhine-Westphalia return program (311-8.03.03.02-147635) and BMBF program Junior Group Consortia in Systems Medicine (01ZX1917B).

