## Supplemental Material for "Deep learning-based segmentation and quantification of podocyte foot process morphology"

### METHODS

**Mouse and human kidney tissue**

All mouse experiments were approved by the State Office of North Rhine-Westphalia, Department of Nature, Environment and Consumer Protection (LANUV NRW, Germany) and were performed in accordance with European, national and institutional guidelines. Mice of 100% C57BL/6N background were used. After anesthesia with Ketamine and Xylazine, mice were euthanized by cardiac perfusion with Hank’s Balanced Salt Solution (HBSS) and fixated as described below.

Control human tissue was collected from patients that were nephrectomized due to renal tumors. Tissue sample was dissected from the non-tumorous pole of the kidney and showed normal histological picture in routine histological examination. All procedures were approved by the Ethics Commission of Cologne and the regional Ethical Committee of Stockholm and conducted in accordance with the declaration of Helsinki. When applicable, patients or the patient’s parents gave informed consent.

### Mouse Model for FSGS

### Mice with two compound-heterozygous point mutations, Pod^R231Q/A286V^, were generated as previously described^10^. Mice were sacrificed at 0 to 20 weeks of age as stated above. Images and ImageJ based analysis data for the Pod^R231Q/A286V^ mice was reused from Butt et al 2020^12^.

### Preparation of kidney tissue for STED/confocal microscopy

Experimental mice were euthanized by decapitation (only newborn mice) or cardiac perfusion with Hanks’ balanced salt solution (HBSS; 5.4 mM KCl, 0.3 mM Na2HPO4, 0.4 mM KH2PO4, 4.2 mM NaHCO3, 137 mM NaCl, 5.6 mM D-glucose, 1.3 mM CaCl2, 0.5 mM MgCl2, 0.6 mM MgSO4) following cardiac blood draw and anesthesia with Ketamine (Zoetis) and Xylazine (Bayer). After median laparotomy, kidneys were removed and fixed in 4% neutral buffered formalin for 2-4 hours at room temperature or overnight at 4°C. After these pieces of kidney were incubated at 4°C in hydrogel solution (HS) (4% v/v acrylamide, 0.25% w/v VA-044 initiator, PBS 1X) over night. The gel was polymerized at 37°C for 3 h, and the presence of oxygen was minimized by filling tubes all the way to the top with HS. Samples were removed from the HS and immersed in clearing solution (CS) (200 mM boric acid, 4% SDS, pH 8.5) at 50°C for 6 h. After that, kidney pieces were cut into 0.3 mm thick slices using a Vibratome. Slices were then incubated at 50°C overnight. Prior to immunolabelling, samples were washed in PBST (0.1% Triton-X in 1X PBS) for 10 min. Alternatively, kidneys were prepared with a recently published fast protocol^7^. Here, samples were fixed for 1-4 hours at room temperature in 4% PFA, and after this kidney pieces were sectioned as described above before incubating sections in CS for 1-2 h at 70°C. Prior to immunolabelling, samples were washed in PBST (0.1% Triton-X in 1X PBS) for 10 min.

### Immunolabelling

### Samples were incubated in a sheep polyclonal antibody to nephrin (R&D systems, AF4269) diluted at 1:50 in PBST (STED microscopy) or 10 mM HEPES pH7.5 with 200 mM NaCL and 10% TritonX-100 (fast protocol) at 37°C for 24 hours (STED microscopy) or 2 hours (fast protocol) with shaking at 500 rpm. For the fast protocol, the antibody was directly conjugated to Alexa-488 using NHS chemistry. After primary antibody incubation, samples were washed in PBST for 5 minutes at 37°C. For the fast protocol, the staining procedure was terminated here. The human sample from a patient with IgAN was stained using the nephrin primary antibody above and a donkey anti-goat secondary antibody conjugated to Alexa-594 (Thermo Fisher, A-11058). For STED microscopy, samples were incubated in a donkey anti-sheep secondary antibody conjugated to Abberior STAR635P (Abberior, 2-0142-007-2, dilution 1:50) at 37°C for 24 hours. The sample from a patient with a mutation in the NPHS1 is lacking nephrin expression in the slit. The slit diaphragm was thus detected by a podocin antibody (Sigma, P0372, 1:100 dilution) and a donkey anti-rabbit secondary antibody conjugated to Alexa-555 (Thermo Fisher, A-31572).

### Mounting

### Samples were incubated in 80% wt/wt fructose (1 mL of dH_2_0 added to 4g of fructose) at 37°C with shaking at 500 rpm for 15 minutes and then placed in a MatTek dish with a cover slip on top (to prevent evaporation) prior to imaging. For the fast protocol, 80% fructose containing 4M Urea was used.

### Imaging

### Images were acquired using a Leica SP8 3X gSTED system using a 100X 1.4 NA objective. A pinhole setting of 0.3 airy units (AU) (as calculated for 594 nm light) was used for the fast protocol unless other stated, for STED imaging the pinhole size was 0.8-1.0 AU.

### Datasets

### For model training and testing we collected a set of 209 STED microscopy images of mouse tissue in varying disease conditions. We incrementally annotated this dataset by first annotating a set of 50 with the macro. We trained an initial model using the 50 image and label pairs and using this model generated predictions for the remaining images. These predictions were then manually corrected using a graphic editor. Our labeling results in 2D segmentation maps with three classes of pixels: background, FP, and SD (Supplementary Figure 1). Each FP in the labeling can be identified as a contiguous pixel blob with the FP label. In all segmentation maps we evened the thickness of the SD to 3 pixels through skeletonization and dilation operations. All images were scaled to the same resolution of 0.0227 microns per pixel. We selected 19 images with varying visual characteristics and set them aside for testing resulting in 90%-10% train-test set split.

### To test the accuracy of inferred morphometric parameters, we used a previously published dataset including 174 healthy and FSGS model mouse tissue samples spanning ages from 0 to 20 weeks old and imaged using STED microscopy^10^. These images were annotated using the macro and the morphometric parameters were also previously quantified.

### Finally, we collected 43 images of human samples representing six different disease conditions (one patient per diagnosis): healthy (7 + 4 images (two patients)), congenital nephrotic syndrome with mutations in the *TRPC6/NPHS2* genes (6 images), minimal change disease (MCD, 4 images), FSGS (12 images), and IgA nephropathy (5 images). These images were generated using the fast protocol including confocal microscopy^7^.

### Segmentation network and training

### We modified U-Net segmentation architecture^21^ by adding two additional convolutional layers after the last U-Net layer. The first convolution produces three output channels and is followed by cross entropy loss. The second convolution produces 16 output channels and is followed by discriminative loss as described previously^17^. This loss forces the 16-dimensional pixel representations to group together, in terms of Euclidian distance, if pixels belong to the same instance and to separate from one another if pixels belong to distinct instances (Supplementary Figure 2). We chose 16 as the smallest dimensionality that results in correct clusters. As full images are too large to fit in the hardware memory, we used 384x384 pixel crops of the original images that were randomly sampled during training. The images were additionally randomly flipped along x- and y-axis and randomly rotated by 0, 90, 180, or 270° during training.

### We used RMSProp optimizer and starting learning rate of 0.05 in training. We additionally used a scheduler that decreased the learning rate by a factor of 0.8 every 100 epochs if no decrease in value of the loss function was observed. We used a batch size of 8 and performed three training iterations of 1,000 epochs each. In each training session the starting learning rate was set at 0.05. We found that this approach resulted in convergence in a better local optimum with each iteration. We performed an additional training iteration on the train set expanded by 55 confocal microscopy images. This training allowed us to adapt the segmentation method to images generated with the fast protocol.

### Inference

### During inference we used windows of size 384x384 pixel that are cropped from the original images with an overlap of 256 pixels between neighboring windows. The overlap allows to better resolve segmentation of FPs that fall on the boundaries of a given window. When merging windows, we take the maximum class label in the overlapping regions. In instance segmentation overlapping instances are merged together.

### To separate individual FP instances, pixel representations are clustered. As the correct number of clusters is unknown, we perform several clusterings for each image window and choose the one that is the best according to a set of criteria. These criteria are the number of distinct clusters that are spatially connected, number of clusters that are outside of a predefined size range, and standard deviation of obtained cluster sizes. Clustering showing minimal sum of ranks in all three criteria is chosen. The clusterings that we test for each image window is defined based on the total area of predicted FP surface in this window. We performed regression of the FP area to the number of FPs in the train set. Numbers of clusters between 0.7 and 1.3 ratio of the expected number of FPs based on this regression are tested as described above. The resulting instance segmentation is additionally postprocessed by removing FP with areas below 0.05 or above 1.5 microns, as well as FPs lying on the image boundaries that are not fully enclosed in an image. We used mini batch k-means clustering with a batch size 1,000 as clustering method. Due to the large number of clusterings we perform during the inference procedure, we parallelized this process across 40 CPU cores.

### To quantify the agreement between predictions and labels we perform an instance matching procedure. In this procedure for each labeled FP, we search for a predicted FP instance that shows the highest area overlap with the given labeled FP. Area of overlap between the matched FPs is quantified as IoU. Predicted instances that do not overlap with any labeled FP are counted as false positives, labeled FPs that have no overlap with predicted instances are counted as false negatives.

### Both training and testing were performed on GPU nodes equipped with four 32 GB NViDIA Tesla GPU cards and two Intel Xeon Gold processors.

### Data labeling using paired microscopy images

### To adapt the network to segment FP in images with lower resolution acquired with the fast protocol, we generated both confocal and STED images of the same field of view. We ran predictions on the STED images using the network trained on STED images as described above. These predictions were then manually corrected and used as labels of the confocal images.

### Automatic ROI assignment

### Macro-based analysis requires ROI to be assigned manually and its boundaries to correspond to capillary area of the glomerulus. Here we approximate this area automatically. First, we find contours of the regions in the semantic segmentation results that do not belong to background. Contours smaller than 0.4μm^2^ are discarded. We next dilate the contour regions for 15 iterations and erode with another 10. We found that through this approach results the contours in close proximity become connected producing a uniform ROI with size approximately matching the one manually assigned.

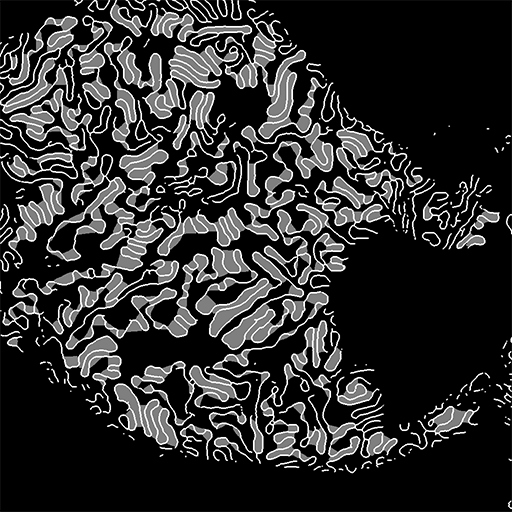

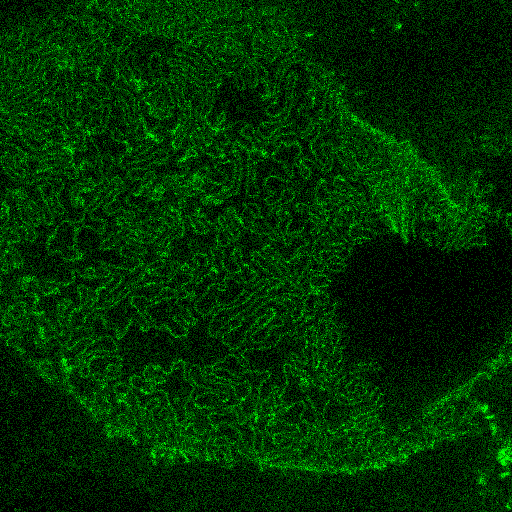

**Supplementary Figure 1.** Labeling. For each original image (left panel) the SD is traced (white marking, right panel), and individual FPs are marked as separate pixel blobs (gray marking, right panel). To facilitate labeling we first trained a segmentation network on a small set of images labeled using the macro. We used this network to perform segmentation in all the images in the train set. These results were manually corrected with the use of a graphic editor. The resulting labeled dataset was used to train the final segmentation model.

Embeddings	Instance segmentation

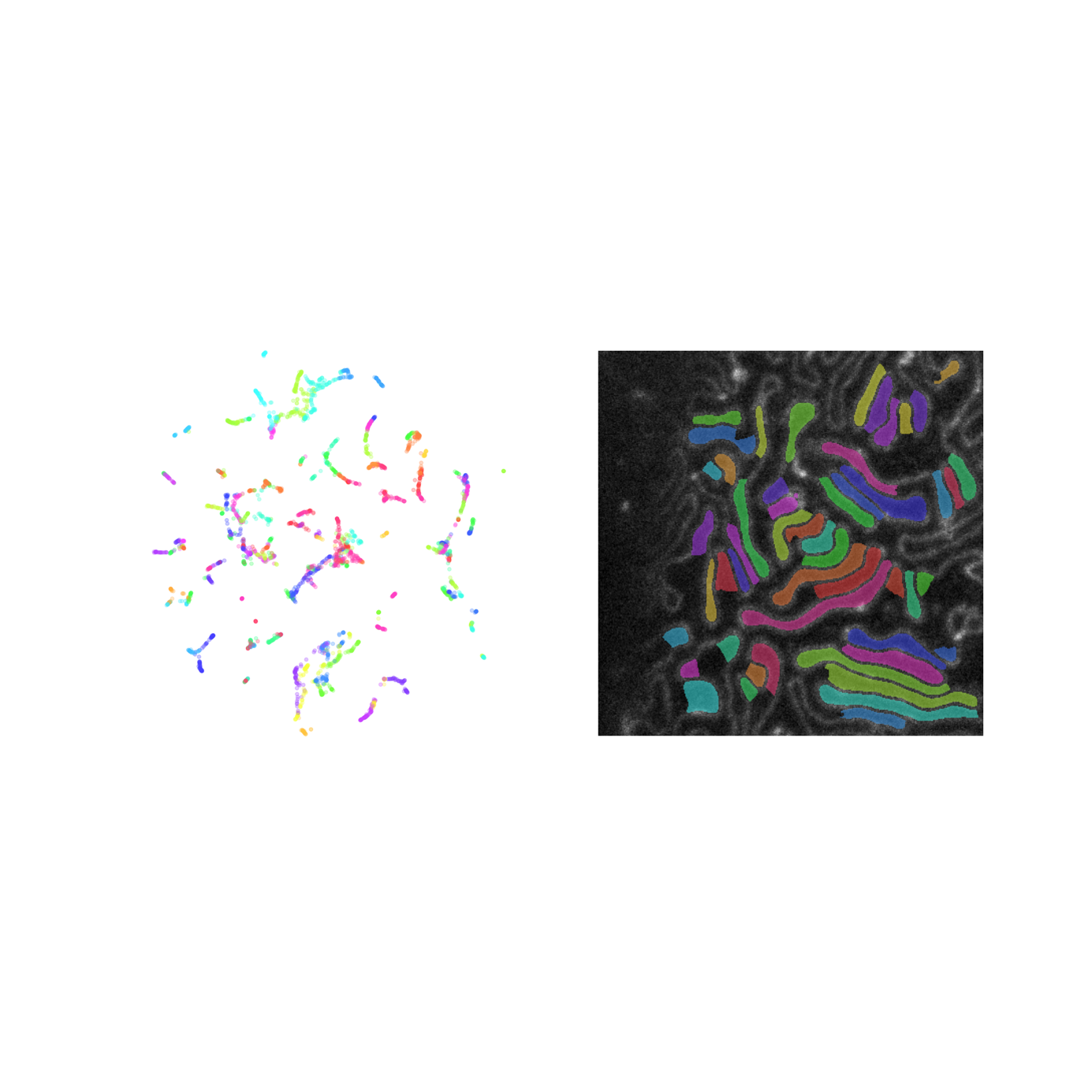

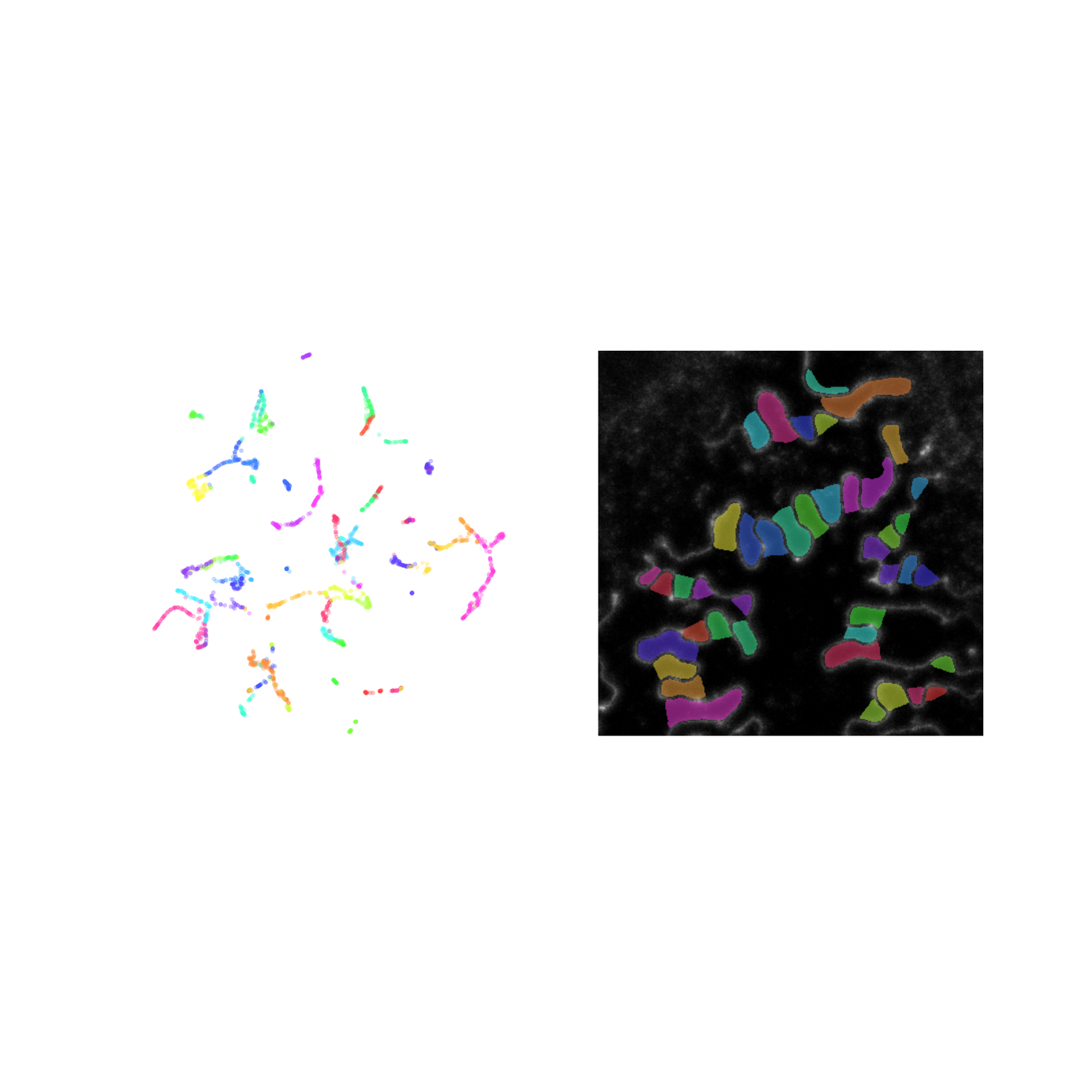

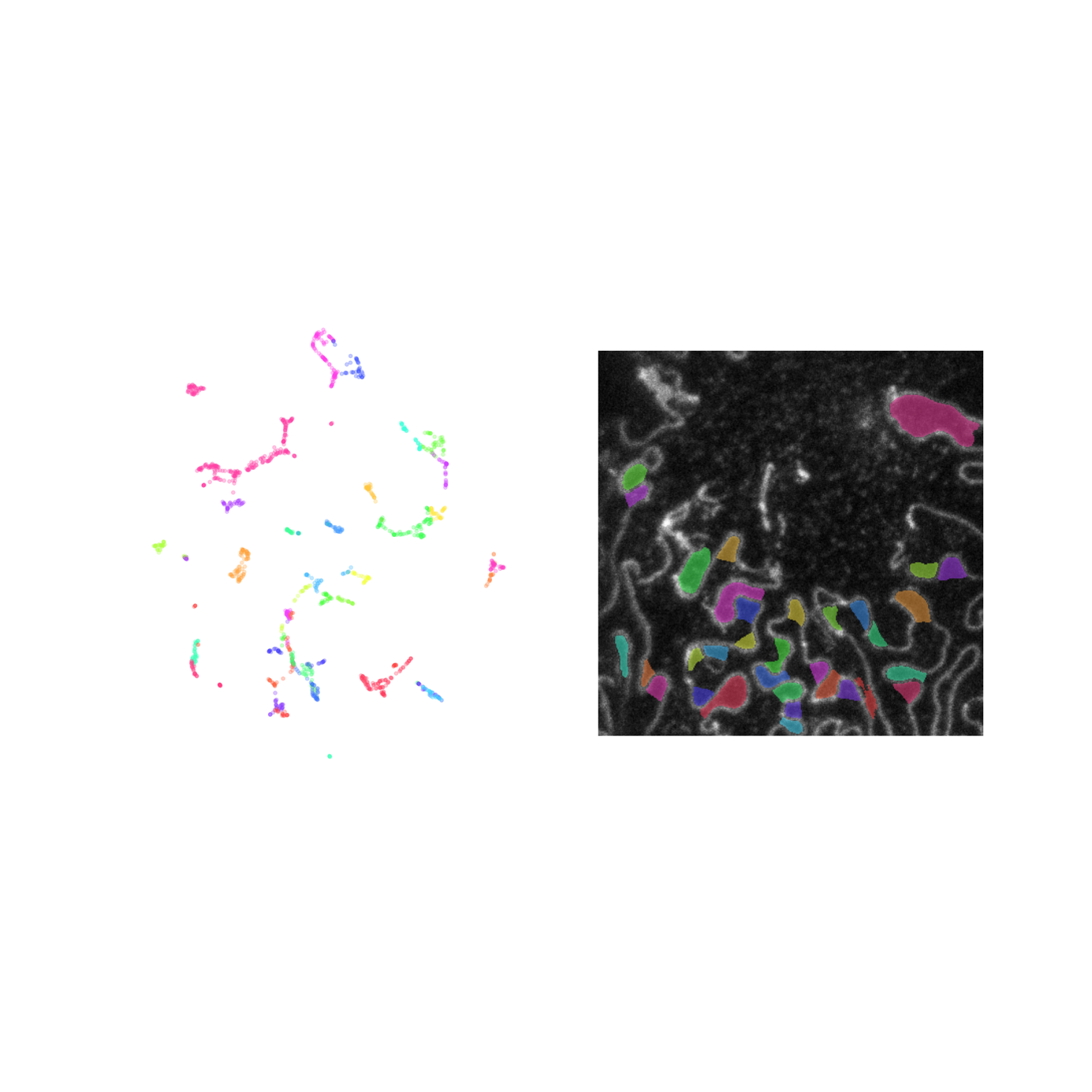

**Supplementary Figure 2.** Assignment of FP instances. Instance segmentation represents predicted FP pixels with 16-dimensional embeddings. These embeddings are clustered, and the resulting clusters correspond to separate FP instances. The panels on the left show a umap visualization of the embeddings of the FP pixels in the respective panels on the right. The colors are matched between the left and right panels. The embeddings of separate FPs group together allowing for their correct clustering and instance assignment.

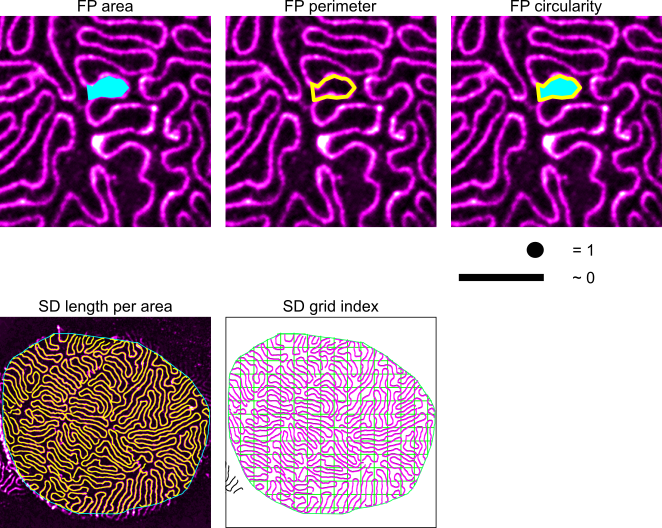

**Supplementary Figure 3.** Quantified FP and SD parameters. FP area = area assigned individual foot process (depicted in cyan). FP perimeter = length of outer boundary of FP (depicted in yellow). FP circularity = dimension-less value expressing how circular a given geometric body is (circularity = 4 * π (area/perimeter²)). A perfect circle has a value of 1, an elongated polygon approximates 0. SD length per area = length of the nephrin signal (yellow) within the ROI (cyan). SD grid index = mean distance between two intersections of the nephrin signal (magenta) with a grid line (green).

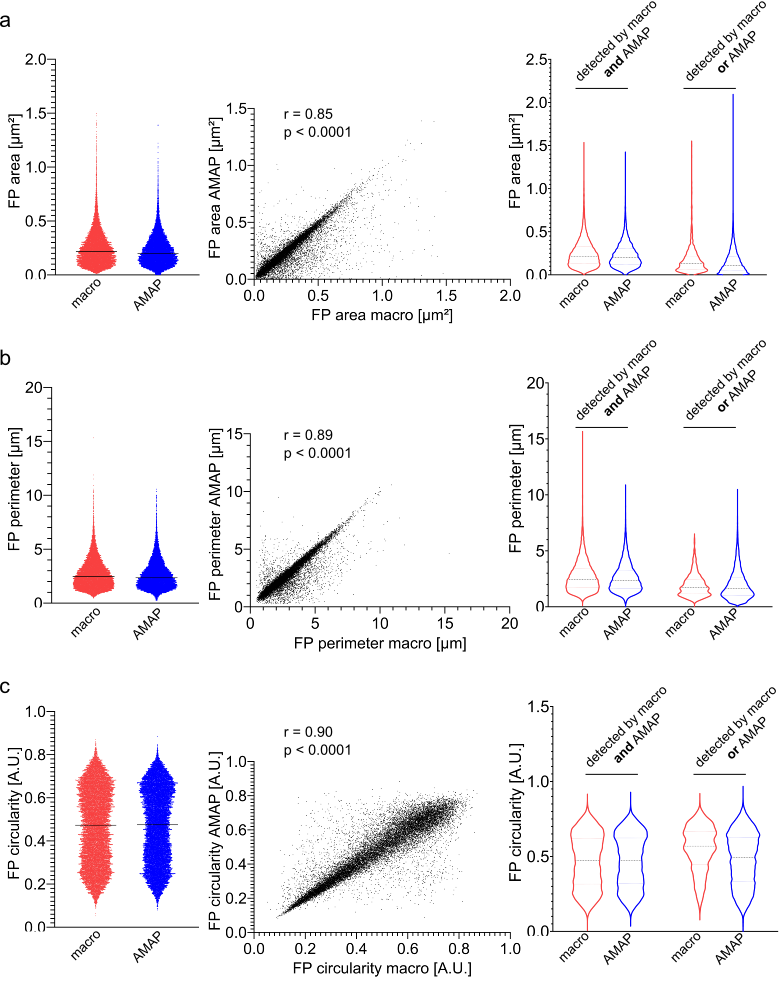
 **Supplementary Figure 4.** Comparison of FP area (a), FP perimeter (b) and FP circularity (c) of FPs which were detected with both the macro (red) and AMAP (blue) (left and middle panels). Each dot represents one FP (n = 18,673 FPs). Horizontal bars represent the median. r = Pearson correlation coefficient, p = p-value. Comparison of FP area (a), FP perimeter (b) and FP circularity (c) of FPs which were detected with both the macro and AMAP or which were only detected either by the macro or AMAP. Dotted lines indicate the median and the quartiles.

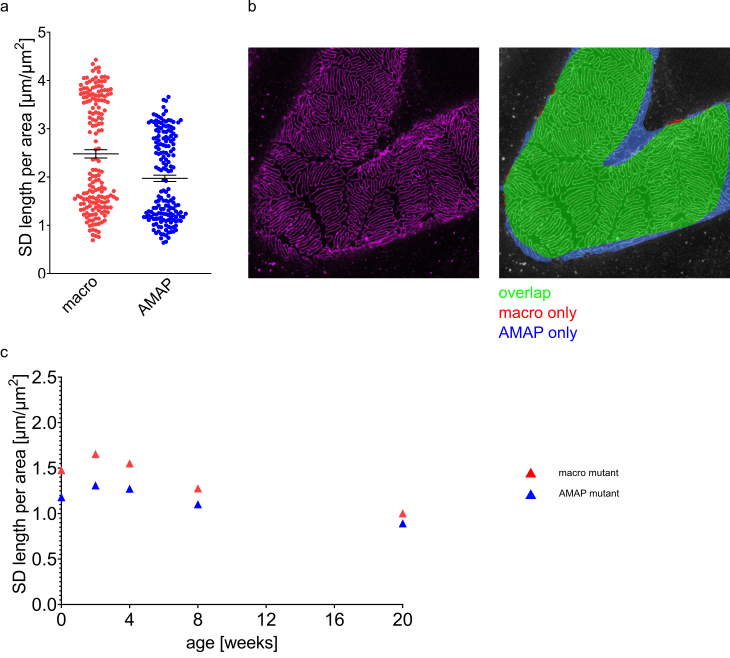

**Supplementary Figure 5.** (a) SD length per area is ~20 % lower in AMAP- assigned ROIs as compared to manually assigned ROIs (left panel). Data are shown as mean ±SEM. (b) Representative original STED image following the immunolabeling of nephrin (left panel) and after the manually and AMAP-assigned ROIs (right panel). Differences and overlap are color coded as indicated in the figure. (c) Mean values of SD length per area (macro in red, AMAP in blue) in mutant mice plotted against their age.

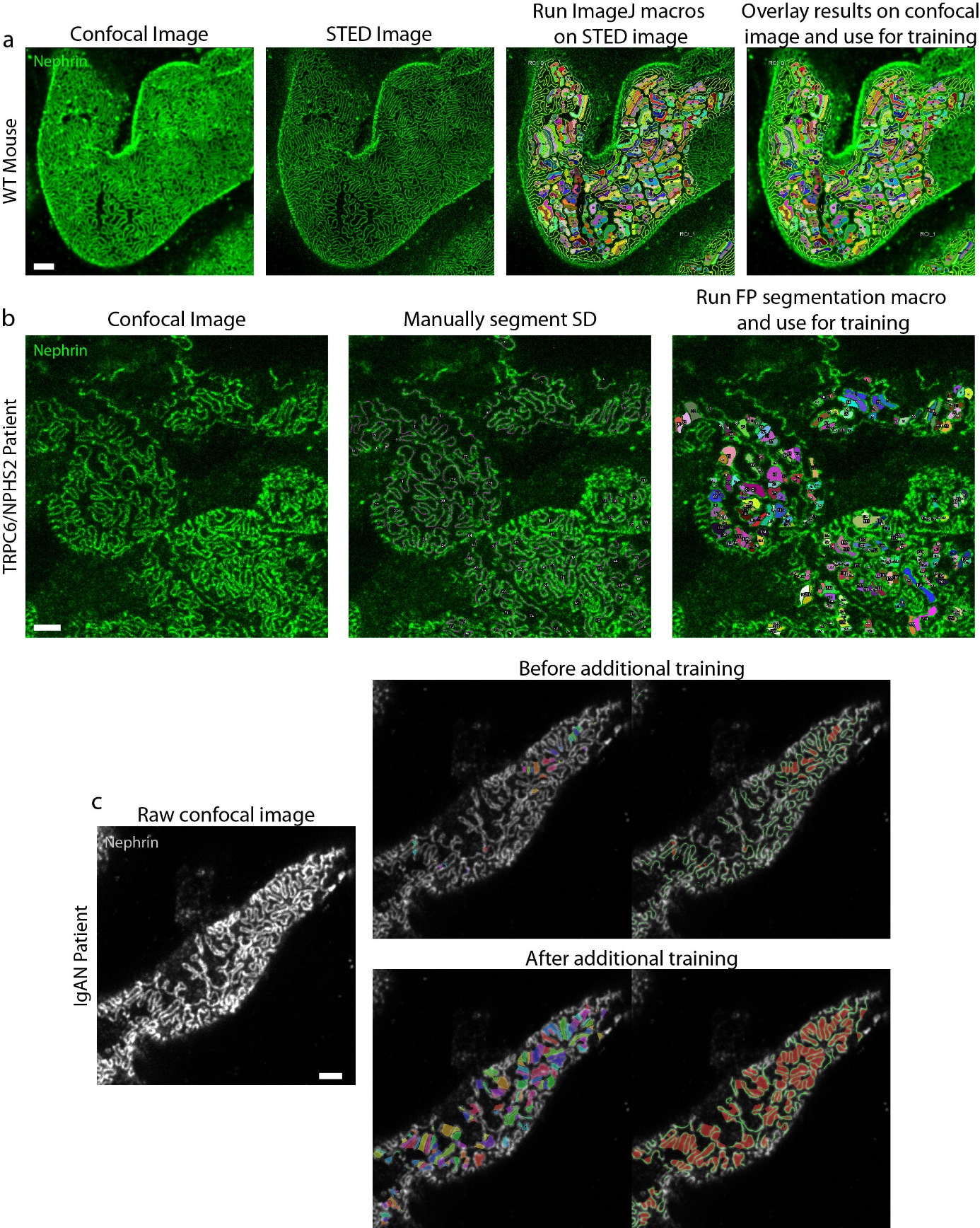

**Supplementary Figure 6.** Generation of training data for the fast protocol images (confocal microscopy). Scale bars 2 µm. (a) One confocal and one STED image was acquired of the same field-of-view. A relatively large confocal pinhole diameter (0.9 airy units) together with a far-red dye (Abberior STAR635P) were applied in order to deliberately decrease the confocal resolution of the training data. The segmentation was applied to the STED images, manually corrected, and the results were then overlaid onto the confocal images which were then used for training of the network. (b) The slit diaphragm in confocal images of the human TRPC6/NPHS2 patient were manually annotated using the freehand line selection tool in ImageJ. Here, images of relatively low quality and contrast were chosen in order to train the network to segment also the lower quality images. (c) Examples of segmentation results before and after training the network with the images in (a-b). The increased segmentation accuracy is evident through visual inspection. For this IgAN patient, a red dye (Atto 594) was used to label nephrin, resulting in lower optical resolution and more challenging segmentation of foot processes as compared to the standard fast protocol where violet/blue/green dyes are used.
